# Warm or bright – temperature and light microhabitat use in insect pollinators

**DOI:** 10.1101/2024.12.14.628486

**Authors:** Océane Bartholomée, Vun Wen Jie, Paul Caplat, Henrik G. Smith, Emily Baird

## Abstract

1. Environmental heterogeneity in forest understories creates microhabitat niches that differ both spatially and temporally in light intensity and temperature. Do animal communities segregate in relation to these niche dimensions and can this be explained by functional traits? Answering these questions is particularly important for insect pollinators as they play a critical role in maintaining flowering plant biodiversity.
2. Bumblebees are essential pollinators of high altitude/latitude ecosystems and are particularly sensitive to climate change. In early spring, they forage on bilberry, a keystone species in heterogeneous habitats – hemi- boreal forests. We capitalized on these conditions to study species-specific selection of foraging niches in relation to abiotic conditions.
3. We combined full-day monitoring of bumblebee communities foraging in bilberry-dominated forests with joint species distribution modelling, which showed that temperature conditioned species occurrence, while light intensity explained species abundance. The inclusion of functional traits did not improve the overall explanatory and predictive power of the models.
4. Our results suggest that temperature acts as a first filter of the local species pool and that species, once present, partition along a light intensity gradient. This study confirms and extends upon previous findings that microhabitat partitioning may act a mechanism underpinning bumblebee coexistence. It highlights the importance of focusing on micro-scales when studying how species interact with their environment, as this could, for example help improve our ability to predict consequences of global changes.

## Introduction

Understanding what determines species’ ecological niches is important for providing insights into the mechanisms that drive community assemblies and can help us predict how community composition will respond to environmental changes (Laughlin et al., 2015; Powney et al., 2014; Suding & Goldstein, 2008). Being able to forecast such responses for insect pollinators is particularly important as they are critical for maintaining the diversity of flowering plants (Ollerton et al., 2011) and support a sizeable proportion of agricultural production (Klein et al., 2007). Over the past decades, bees – the most important group of insect pollinators – have been declining due to anthropogenic disturbances (Potts et al., 2010; Wagner, 2020) such as habitat loss (Winfree et al., 2009), pathogens (Cameron et al., 2016; Meeus et al., 2011), and climate change (Kerr et al., 2015). While our knowledge about how bee communities respond to global change is increasing, our understanding of the underlying mechanisms is still poor – particularly with respect to how species respond to variations in abiotic conditions at small spatio-temporal scales.

The coexistence of many bee species has long been a focus of research on niche separation. For instance, within a specific habitat up to 16 species of bumblebees can coexist (Goulson et al., 2008). This extensive coexistence has been suggested to be facilitated by differences in tongue length, which could limit inter-species competition by facilitating exploitation of distinct flower species (Heinrich, 1976; Inouye, 1978; Ranta & Vepsäläinen, 1981; Sponsler et al., 2022). Differences in tongue length do not directly explain bumblebee species coexistence in hemi-boreal forests, where the flowering resource is often homogenous due to dominance by a single or a few species. However, in these environments, abiotic factors often vary at small scales because of the shading by vegetation is dynamic and changes with the position of the sun. This creates a small-scale heterogeneity in light (light intensity and spectra, (Théry, 2001)) and micro-climatic conditions (Haesen et al., 2021), which could potentially drive niche partitioning in ectotherms (Woods et al., 2015).

Following the ‘trait-based environmental filtering’ hypothesis (Vellend & Agrawal, 2010), we expect to find mobile species in (micro)habitats where their functional traits provide optimal performance – such as foraging success – under specific local environmental conditions (Green et al., 2022; McGill et al., 2006). For example, for insect pollinators, tongue length reflect their ability to access nectar in flower with different corolla depths (Ranta & Lundberg, 1980). However, trait-environment relationships can be more complex, with some traits having several functions. For example, bee body size affects their mobility (Greenleaf et al., 2007) and their temperature sensitivity as, according to Bergmann’s rule, larger species have higher cold tolerance (Bishop & Armbruster, 1999). Moreover, compound eye characteristics can affect insect activity periods, *e.g.* diurnal, crepuscular and nocturnal bees have different visual traits (Dorey et al., 2020) but they can also affect habitat preference, as has been shown in damselflies (Scales & Butler, 2016), *Drosophila* (Keesey et al., 2020) and butterflies (Wainwright & Montgomery, 2022). Studies focusing on trait-environment relationships can get complicated due to correlations between functional traits (Estrada et al., 2016; Fitzgerald et al., 2022). For example, in bees, body and eye sizes are allometrically related (Jander & Jander, 2002), with larger bees – trait associated with higher cold tolerance (Peters et al., 2016) – having larger eyes and a higher light sensitivity (Kaputjanski et al., 2007; Streinzer et al., 2016). An ideal trait for investigating how species respond to varying light intensity is therefore the eye parameter, which reflects the trade-off between light sensitivity – *i.e.* dim light vision – and visual resolution, that can vary independently from both body size and eye size (Jander & Jander, 2002; Taylor et al., 2019; Tichit et al., in preparation). In bumblebees, the eye parameter has previously been related to habitat use (Tichit et al., in preparation) and light microhabitat partitioning (Bartholomée et al., 2023), such that species with higher eye parameters (higher light sensitivity) have microhabitat niches that include dimmer light conditions. Functional traits such as body size and eye parameter can therefore be useful for exploring their role in how insect communities partition across both temperature and light niches.

In this study, we aim to improve the understanding of how variation in temperature and light intensity conditions at small spatio-temporal scales affect the community composition of insect pollinators. We do this by focusing on bumblebees in Nordic hemi-boreal forests that, in early spring, forage on bilberry (*Vaccinium myrtillus*), which dominates the understory and constitutes a primary food source for at least ten bumblebee species (Andresen, 2019; Bartholomée et al., 2024; Moquet et al., 2017). A recent study suggested that species coexistence in these habitats could be explained by niche-separation based on abiotic conditions (Bartholomée et al., 2023). It focused on the relationship between light microhabitats and eye traits and found that bumblebee communities foraging on bilberry segregate by using different light microhabitats and that a species’ light microhabitat niche can be explained by variations in visual abilities (Bartholomée et al., 2023). However, this earlier study was limited to observations made in the afternoon, when there was little variation in temperature. It thus remains unclear how variations in other environmental variables, such as temperature, affect how bumblebees use microhabitats and whether this is supported by their functional traits. We focused our analyses on bumblebee queens to avoid any potential caste specific differences. We expect bumblebee species to partition along light and temperature gradients, with those active in the dimmer and colder periods of the day to have larger bodies, and higher eye parameters, as these functional traits would facilitate foraging under these conditions.

## Methods

### Field sites

The study was carried out in two selected plots of forested land (∼150 m x 50 m) located near Tovetorp Zoological Research Station in Södermanland County, Sweden (50 m.a.s.l, 58°56’N, 17°08’E). The plots contained hemi- boreal forests dominated by Norway spruce (*Picea abies* (L.) H.Karst.) and Scots pine (*Pinus sylvestris* L.), with an understory dominated by the European bilberry (*Vaccinium myrtillus* L.). The two plots had contrasting openness. The ‘closed’ plot had a dense canopy due to a higher number of young Norway spruces, while the ‘open’ plot, was brighter and its understory was interspersed by lingonberry (*V. vitis-idaea* L.).

### Observation transects and environmental variables

A fixed loop transect corresponding to a 15 min walk at a slow speed (∼300 m) was set up within each plot (Table 1). Bumblebee monitoring along these transects was performed at the peak of bilberry blooming, between the 12^th^ and 22^nd^ of May 2022. We monitored bumblebee communities on both plots from 30 min before sunrise to 30 min after sunset during seven days (12/05: sunrise at 04:29, sunset at 21:08; 22/05: sunrise at 04:09, sunset at 21:29). To assess the effects of rapidly changing light intensity, we performed observations from 30 min before to 30 min after both sunrise and sunset for one of the two plots for each monitoring day. During this 1 h monitoring period, we walked the fixed transect four times. Between sunrise and sunset, transects in both plots were walked for 15 min every hour – from either 60 min after sunrise (for the plot selected for sunrise/sunset), or 90 min after sunrise (for the other plot). When bumblebee activity was very high, the transects could not be walked in their entirety as recording observations took more time. To record bees over a wide range of light intensities, we included observations in all weather conditions except rain.

**Table 1:**
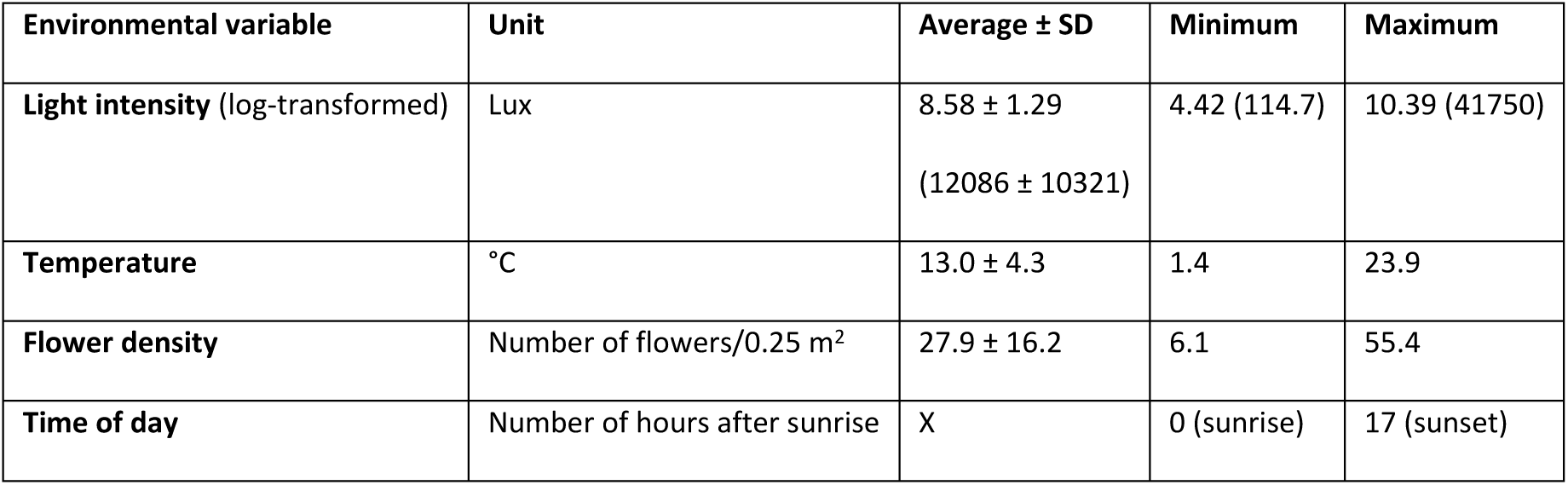
Environmental variables monitored in the study at the transect level (226 transects): average, standard deviation (SD), minimum and maximum X indicates that the average is not meaningful.

During each transect, all observed bumblebees were noted and identified to species and caste when possible, based on a Swedish bumblebee guide (Söderström et al., 2021). Bumblebee species that could not be reliably identified in the field due to their similar appearance (*Bombus terrestris* (Linnaeus, 1758)*, B. lucorum* (Linnaeus, 1761)*, B. magnus* (Vogt, 1911), and *B. cryptarum* (Fabricius, 1775)) were counted as *B. terrestris* complex (BTC) (Carolan et al., 2012). Similarly, the cuckoo species *B. sylvestris* (Lepeletier, 1832) and *B. bohemicus* (Seidl, 1837) were grouped, as they were also difficult to distinguish reliably. For each individual observation, we recorded the (i) the day, (ii) the time (coded as integer from 0 to 17, representing the number of hours after sunrise, with sunrise being 0 and sunset 17), (iii) light intensity (measure span between 0 and 200 klux, accuracy: ± 4 % rdg ± 0.5 % f.s. (<10,000 lux)) by maintaining the light sensor parallel with the sky, and (iv) temperature (measure span between -40°C and 70°C, <± 1.0°C) (for both variables: DEM900 Environment meter, Velleman Group, Belgium) (Table 1). For each of the two sites, we measured flower density daily by counting the number of bilberry flowers in eight randomly located 0.5 m x 0.5 m quadrats (0.25 m^2^) (Table 1). The counts were averaged over the eight replicates.

### Bumblebee functional traits

To explain the use of light and temperature microhabitats by bumblebee species/caste combinations, we focused on two traits. We included the intertegular distance (ITD, *i.e.,* the distance between the forewing bases) as a measure of body size that reflects temperature sensitivity with larger bees having a higher cold tolerance (Bishop & Armbruster, 1999; Gérard et al., 2018). We use this trait to represent thermal adaptations here because, while bumblebee body size does reflect other life-history and physiological characteristics (Fitzgerald et al., 2022), we do not consider them relevant for the foraging context studied here. In addition, as previous work on similar bumblebee communities in a similar habitat found no correlation between body size and light niche (Bartholomée et al., 2023), we also consider these factors to be independent, although we acknowledge that previous work has suggested that, generally, larger bees able to fly in dimmer light conditions (Kaputjanski et al., 2007). Body size measurements were made on 10 queens/species from the collections of the Biological Museum Lund University (Romain Carrié, unpublished data). For the cuckoo species, we used measurements for *B. sylvestris* from Kendall et al. (2020) (Table S1). For the BTC, we averaged the ITD for *B. terrestris* and *B. lucorum.* Eye parameter was used as the visual trait as it has previously been related to light microhabitat associations in bumblebees (Bartholomée et al., 2023). The eye parameter is defined as the product between the facet diameter (μm) and the angle between two adjacent facets (radian) and is a measure of the trade-off between the investment in visual sensitivity and resolution – resulting, respectively, in bigger facets and smaller inter-facet angles. The eye parameter measures were taken from Tichit et al. (in preparation) (Table S1). As in Bartholomée et al. (2023), if the eye parameter values for a queen were missing, we used values taken from workers of the same species. This can be justified due to the low intra-specific variation in this trait (Tichit et al., in preparation), *i.e.* for *B. soroeensis* (Fabricius, 1776), *B. hypnorum* (Linnaeus, 1758) and *B. lapidarius* (Linnaeus, 1758). For the *B. terrestris* complex, values from *B. lucorum* and *B. terrestris* were averaged (Table S1). For the cuckoo group, we used the measurements available for *B. bohemicus*, as measurements for the other species of the group were unavailable (Table S1). The sample size for the eye parameter values is low, which is explained by the complexity and the time required for the calculation of this trait (Jezeera et al., 2022; Taylor et al., 2019). Nonetheless, we believe that our data are useful, because this trait is highly species-specific and only weakly related to body size (Tichit et al., in preparation).

### Statistical analyses

All analyses were performed using R version 4.3.1 (RCoreTeam, 2022) and associated packages (shown in italics). We analysed data at the transect level, *i.e.,* data were agglomerated for each transect, such that microhabitat light and temperature conditions where those experienced in average by the bumblebees during their activity at the time of observation. Light intensity and temperature were calculated as the average light intensity and temperature taken for each identified bumblebee observation during each transect walk (cf. Bartholomée et al., 2023). As workers represented only 3.4% of the observations, we excluded them from the data set to avoid any potential caste-related biases. Light intensity values were log-transformed before analyses because they varied over several orders of magnitude.

We used Joint Species Distribution Modelling (Ovaskainen et al., 2017) to describe how the variation in bumblebee community composition is related to both environmental (Tarafdar et al., 2021) and species traits. More specifically, we applied Hierarchical Modelling of Species Communities (HMSC) as implemented in *Hmsc* (v 3.0, Tikhonov et al. (2020)). In these analyses, we considered each transect walk as one sampling unit and used the model’s default priors (Ovaskainen & Abrego, 2020).

To handle the zero-inflated data, we chose a hurdle approach that comprises two models (Bosco et al., 2023; Pena et al., 2023): a first model for species occurrence where the data was truncated to absence (0) and presence (1) values with a ‘probit’ (binomial) distribution, and a second model for species abundance conditional on their presence with a log-normal distribution. For the latter model, species absences were considered as missing values, while abundance data were log-transformed. The hurdle approach allowed us to separately assess the drivers of species occurrence and the drivers of the abundance of the species that were present. To ensure comparability between different transect lengths and as species richness was not significantly different between the observations taken at sunrise/sunset (for 1h) and during the rest of the day (15 min of observation; linear regression, *F*1, 224 = 2.30, *P* = 0.13), we converted community observations to the equivalent of 1 h observation time by multiplying by 4 the observations that were taken over 15 min (*i.e.,* those taken outside sunrise and sunset). We removed species with less than 10 individual observations (*B. hortorum* (Linnaeus, 1761)).

In all six models (3 model structures x 2 distributions), we included three crossed random effects to account for: (i) plot identity to account for unmeasured environmental variation between the two sampling plots, (ii) date of monitoring to account for repetitive measurements, both coded as unstructured random effects and (iii) a temporally structured random effect for time of day to account for potential temporal correlation (cf. Wells et al., 2021). These random effects accounted for variation over time (within plots) and space (between plots) – as intra-transect variations were not included because of the light-temperature conditions averaging per transect walk.

For both probit and log-normal data we ran three models. First, we built a null (intercept only) model – M0 – without predictors, to assess whether the random effects were influencing bumblebee species. To test if there is micro-habitat partitioning, we ran M1 where we included three measured environmental variables as predictors (significantly related to community composition in NMDS analysis, Figure S1): (P1) light intensity (lux, log-transformed), (P2) temperature (°C), and (P3) flower density (number of flowers/0.25 m^2^). For both light intensity and temperature, we included second-order polynomial functions (Bosco et al., 2023) as we observed non-linear responses in the quadratic trend surfaces overlaid on NMDS analyses (Figure S1). To assess if the observed patterns were supported by functional traits, we built M2, by including the two functional traits to M1: (T1) intertegular distance (ITD, mm), (T2) eye parameter (μm.rad).

For all models, we used four Markov chain Monte Carlo (MCMC) chains with each 1 500 000 iterations, out of which we ignored a transient of 500 000 iterations to sample the posterior distribution. The remaining iterations were thinned by 1000, leading to 1000 posterior samples per chain, so 4000 posterior samples in total. To check the MCMC convergence, as described in Tikhonov et al. (2020), we considered both the effective sample size of MCMC samples and the potential scale reduction factor through the Gelman-Rubin convergence statistic (Gelman & Rubin, 1992), which indicates convergence by a value close to 1 (Bosco et al., 2023).

We compared the three models with their explanatory power and their predictive power after a two-fold cross- validation (Barrero et al., 2023; Pena et al., 2023). For the occurrence models, we used the Tjur’s R^2^, which is a measure of the difference between occurrence probability in sampling transects where species are present or absent. For the models of abundance conditional on presence, we used the standard coefficient of determination R^2^.

To quantify the proportion of the explained variance of species occurrence and abundance explained by the different environmental variables and random effects, we used variance partitioning (Ovaskainen et al., 2017). For M2, we quantified the proportion of species responses to environmental variables explained by traits. In addition, we assessed the posterior distributions of the models’ parameters (*i.e.,* the level of support from the MCMC sampling) to assess how individual species respond to environmental variables and how traits relate to these environmental variables (cf. Wells et al., 2021).

## Results

We performed 67 h of transects and counted 3877 bumblebee individuals coming from eight species/group of species (Table 2).

**Table 2:** Community composition of the 3877 individuals counted during 67 h of transect walks (including 14 h around sunrise and sunset) in individual light intensities between 2.6 lux and 106000 lux (0.9 and 11.6 in log-lux) and temperatures between 1.0 °C and 26.5 °C. X indicates that counts for that caste was impossible (as cuckoo species do not have workers) or unavailable as non-identified (for *Bombus* spp.).

| Species | Total number (% of observations) | Number of queens | Number of workers |
| --- | --- | --- | --- |
| BTC* | 2641 (68.1%) | 2600 | 41 |
| *B. pratorum* (Linnaeus, 1761) | 307 (7.9%) | 238 | 69 |
| *B. pascuorum* (Scopoli, 1763) | 228 (5.9%) | 218 | 11 |
| *B. soroeensis* (Fabricius, 1776) | 215 (5.5%) | 209 | 6 |
| *B. lapidarius* (Linnaeus, 1758) | 198 (5.1%) | 198 | 0 |
| Cuckoo** | 155 (4.0%) | 155 | X |
| *B. hypnorum* (Linnaeus, 1758) | 51 (1.3%) | 46 | 5 |
| *B. hortorum* (Linnaeus, 1761) | 7 (0.2%) | 7 | 0 |
| *Bombus* spp. | 74 (1.9%) | X | X |

\**Bombus terrestris* complex
\*\**B. sylvestris* and *B. bohemicus*

We used joint species modelling to estimate species responses to environmental drivers and if this was modulated by functional morphological traits. The convergences of the MCMC for the six models - both the occurrence (probit) and the conditional abundance models for the models M0, M1 and M2 – were satisfactory as the effective sample sizes of both Beta and Gamma parameters were close to the real number of samples (4000) and the potential scale reduction factors of these parameters were generally close to one (Table S2).

The random effects explained between 1 and 25% of species occurrences and between 0.18 and 70% of species conditional abundance, highlighting the role of random effects on explaining species responses – with the relative importance of the random effects depending on the species (Table S3). The higher explanatory and predictive power of M1 and M2 compared to M0 emphasizes the role of the environmental variables included in the models. For species occurrence, M1 and M2 were similarly supported in terms of explanatory and predictive powers (highest Tjur’s R^2^) (Table 3). For the conditional abundance, M1 and M2 had identical explanatory power (R^2^, Table 3), although M1 had a higher predictive power in the cross-validation. As including functional traits in the models did not improve their fit nor predictive powers, we selected M1 over M2. Therefore, we subsequently focus on the results of M1. Results of M0 and M2 are detailed in the Supplementary material (respectively Table S3 and Tables S6-S11).

**Table 3:** Comparison of the model fit for Models 0, 1 and 2 for both occurrence and conditional abundance: explanatory and predictive powers as estimated by the Tjur’s R^2^ for occurrence models and R^2^ for conditional abundance models.

| | Explanatory power<br>Average (± SD) [min; max] | Predictive power<br>Average (± SD) [min; max] |
| --- | --- | --- |
| **Occurrence model** |  |  |
| Model 0 (intercept only) | 0.09 (± 0.09) [0.01; 0.25] | 0.05 (± 0.06) [-0.004; 0.16] |
| Model 1 (environment) | 0.15 (± 0.08) [0.05; 0.27] | 0.10 (± 0.08) [-0.007; 0.24] |
| Model 2 (environment + traits) | 0.16 (± 0.08) [0.07; 0.29] | 0.10 (± 0.09) [-0.001; 0.26] |
| **Conditional abundance model** |  |  |
| Model 0 (intercept only) | 0.38 (± 0.19) [0.18; 0.70] | 0.12 (± 0.14) [0.004; 0.41] |
| Model 1 (environment) | 0.40 (± 0.22) [0.20; 0.75] | 0.27 (± 0.20) [0.04; 0.56] |
| Model 2 (environment + traits) | 0.40 (± 0.22) [0.20; 0.75] | 0.23 (± 0.22) [0.01; 0.56] |

The occurrence model had lower explanatory and predictive powers than the conditional abundance model, as they respectively explained (predicted) between 5% (-0.007%) and 27% (24%) of the observed variation in species occurrence, and between 20% (0.04%) and 75% (0.56%) of the observed variation in species abundance (Table 2, Figure 1A, 1B). The explanatory power between species was highly heterogeneous and lower for BTC, *B. lapidarius* and *B. soroeensis* for occurrence, and lower for *B. soroeensis* and *B. hypnorum* for conditional abundance. The discriminatory power for BTC in the occurrence model needs to be considered with care due to the lack variability in its presence data – this species group was observed in all transects.

**FIGURE 1:**
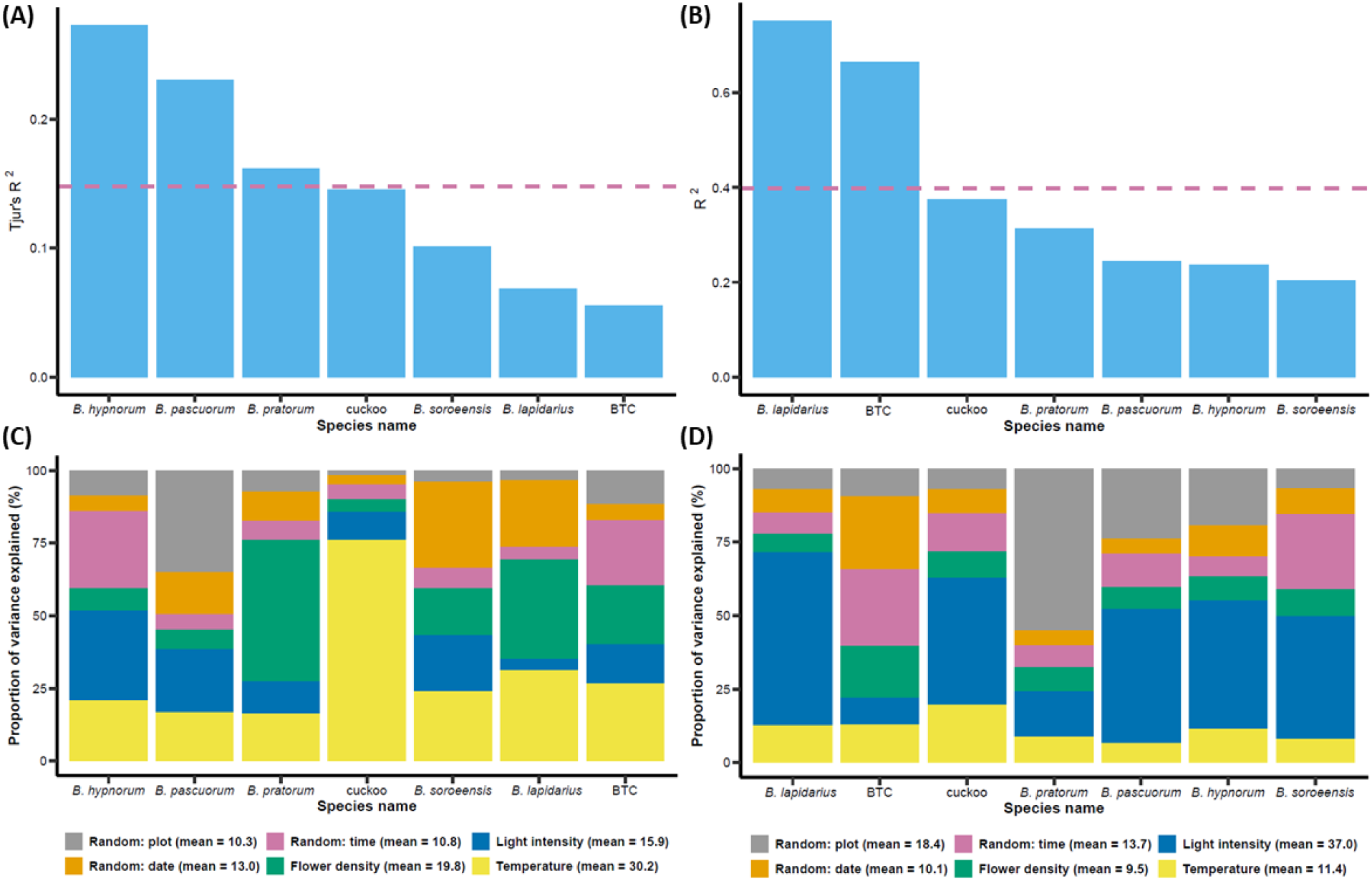
Variance in bumblebee species occurrence and conditional abundance. (A-B) Part of the total observed variance explained by the occurrence (Tjur’s R^2^, A) and conditional abundance (R^2^, B) M1 for the different species/groups. (C-D) The purple dashed lines indicate the average explained variance. Partitioning of this explained variance for the occurrence M1 (C) and the conditional abundance M1 (D). The main environmental variables light intensity (in blue) and temperature (in yellow) indicate the variance due to species’ microhabitat preferences, while flower density (green) shows the differences due to changes in flower availability. The rest of the variance explained is partitioned between the three random effects: 1) plot identity (open or closed) (in grey), which accounts for the unaccounted natural variation between the sampling plots, 2) monitoring date (in orange) explaining inter-day variation (*e.g.,* due to varying meteorological conditions) and 3) time of day (in purple), which accounts for the potential temporal auto-correlation between transects monitored at successive times of the day. The average proportion of explained variance for each variable is specified between brackets in the legend. BTC stands for *B. terrestris* complex.

Variance partitioning showed that light intensity and temperature were the most important explanatory variables of the explained variance. For species occurrence and abundance, light intensity explained 26 % and 30 % of the explained variance, while temperature explained 32 % and 17 % of it, respectively (Figure 1C, 1D). Thus, temperature was the primary driver of species occurrence (30 % of the explained variance across species) and light intensity was the primary driver of the abundance of the present species (37 % of the explained variance across species). The effects of temperature and light intensity differed greatly between species. Light intensity explained 4-31 % of the variance explained of species occurrence and 9-59 % of the variance explained of species abundance. Temperature explained 16-76 % of the variance explained of species occurrence and 7-20 % of the variance explained species abundance (Figure 1C, 1D). Temperature was particularly important for the occurrence of cuckoo species and *B. lapidarius*, while light intensity was particularly important for the conditional abundance of *B. lapidarius*, *B. pascuorum, B. hypnorum* and cuckoo species. Flower density explained 4-49 % of variance explained of species occurrence and 6-18 % of explained variance of species conditional abundance. Both spatial and temporal random effects were more important in the abundance model (42 % of explained variance) than for the species occurrence model (34 % of explained variance).

A posterior probability ≥ 95 % was found between species and the environmental variables. The occurrence probability varied both with species and with environmental variables. The occurrence of all species, besides BTC was positively associated with the temperature linear term, while the occurrence of *B. lapidarius*, *B. pratorum*, *B. soroeensis* and cuckoo species was positively associated with the second-order term, indicating a lower occurrence probability at intermediate values (Figure 2A). The probability of presence of *B. lapidarius* and *B. soroeensis* increased with flower density, while presence for *B. pascuorum* and *B. pratorum* decreased with flower density (Table S4). Light intensity did not influence the occurrence of any species (Figure 2C). For conditional abundance, all species responded positively to the linear effect of light intensity and negatively to its second-order term, indicating an abundance peaking at intermediate light intensity values (Figure 2D) when the abundance of *B. hypnorum* and *B. terrestris* complex peaked for intermediate temperatures (Figure 2B). Cuckoo species, *B. hypnorum* and BTC abundance increased with flower density, while the abundance of *B. pratorum* decreased with flower density (Table S5). In both M2, no trait showed a posterior probability > 95%, which might be due to the relatively low number of species, and validates the selection of M1 (Tables S10 and S11).

**FIGURE 2.**
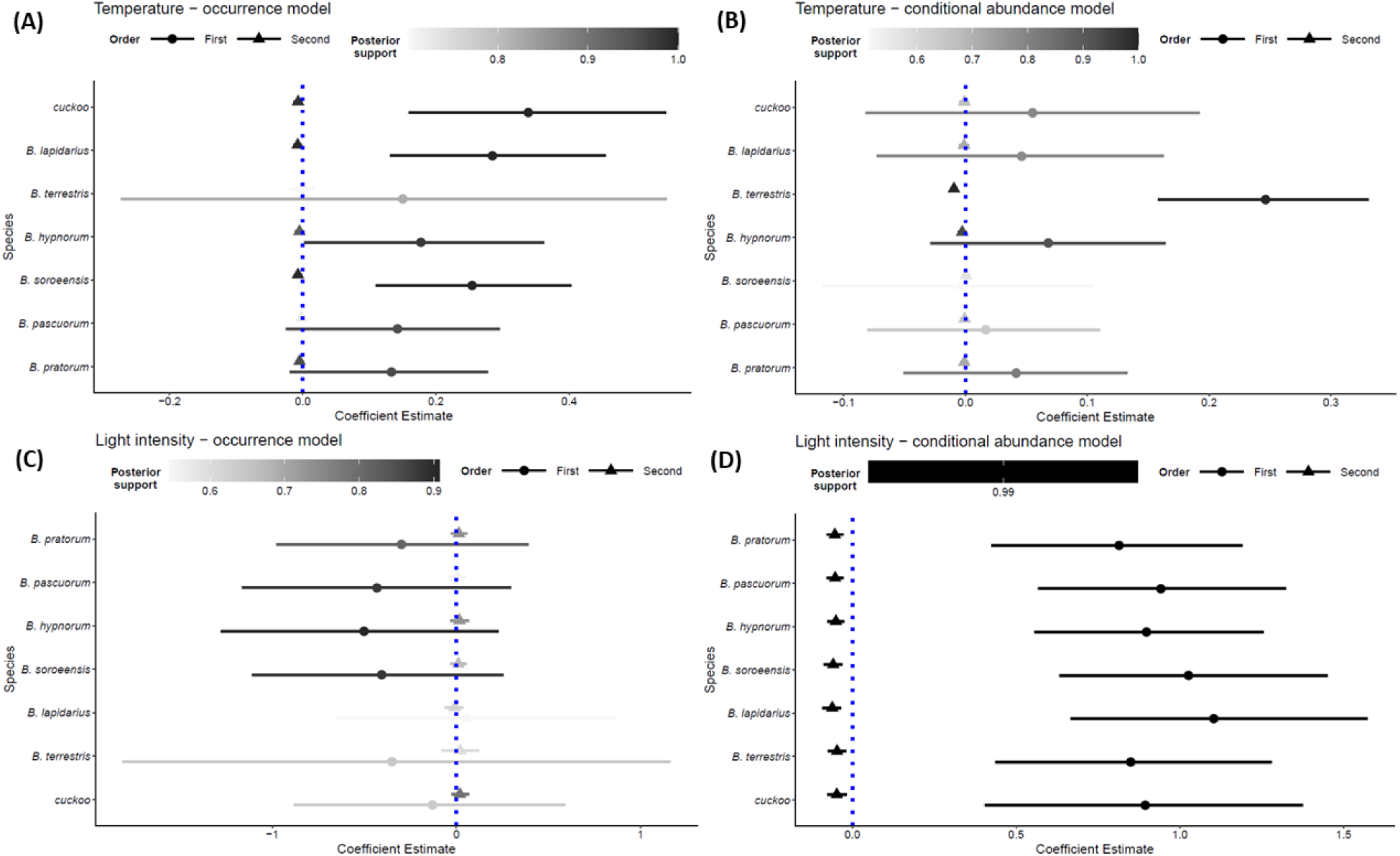
Species-level responses to temperature and light intensity. Responses of species occurrence (A) and conditional abundance (B) to temperature based on the posterior probability in the HMSC M1. Responses of species occurrence (C) and conditional abundance (D) to light intensity based on the posterior probability in the HMSC M1. The central dots are the average Beta parameters and the lines represent the 95% credible intervals. If the lines overlap with the blue dotted line of the 0 value, it means that the environmental variable had no effect on the species response. Circles and triangles correspond to the first and second order terms, respectively, in the environmental equations. The grey gradient scales represent the posterior support of the *Beta* parameters. BTC stands for *B. terrestris* complex.

## Discussion

By combining insect and microhabitat monitoring with joint species distribution modelling, this study represents a novel approach to studying insect pollinator community microhabitat use. We used a study system that is uniquely suitable for this purpose, with a phylogenetically well-resolved diverse pollinator group foraging on a single flower resource, growing in a habitat with heterogeneous microhabitats in terms of light intensity and temperature. We showed that bumblebee species foraging on bilberry were filtered by temperature and that, when present, species partitioned along a light intensity gradient. Traits related to temperature sensitivity (body size, ITD) and light sensitivity (eye parameter) explained between 20 % and 60 % of the variation explained of the species response to variations in temperature and light intensity – as shown by both M2 (Table S6). However, the inclusion of these traits did not improve the explanatory and predictive powers of the models, which might be explained by the low number of species (seven) and the lack of sufficiently large inter-specific variation. Overall, our investigation of how microhabitat variations contribute to shaping bumblebee community dynamics, and highlights the importance of canopy gaps for generating the heterogeneous light and temperature conditions (Muscolo et al., 2014) that facilitate species coexistence. This is important as such canopy gaps are becoming scarcer in modern forestry management regimes (Hamřík et al., 2023).

The results of this study provide good evidence that bumblebee community composition is affected by multiple environmental variables. Our day-long measurements allowed us to decouple light intensity and temperature, which enabled us to further advance the findings of Bartholomée et al. (2023), who found that variation in light intensity microhabitats was a significant driver of bumblebee community composition foraging on bilberry. By covering a broader range of temperatures (1.5-24.0°C compared to 10.0-28.0°C in Bartholomée et al. (2023)), we could determine that, in addition to light intensity, temperature is also a central factor shaping bumblebee communities and that these variables are not necessarily correlated in forest habitats. The effects of time of observation seen in this study is consistent with the differences in bumblebee communities that Bartholomée et al. (2023) observed between noon, afternoon and evening monitoring sessions. Our results also reveal that flower density (not measured in Bartholomée et al. 2023) is an important variable that affects bumblebee community composition (Gómez-Martínez et al., 2020). Taken together, our findings suggest that monitoring insect pollinator communities across the day and over a broader range of temperatures can give a more detailed understanding of how these communities exploit variations in microhabitats. Our results also highlight the importance of considering the relevant scale when measuring micro-climatic conditions in the context of insect studies, as these might change quickly over time and space, and the scale at which the insects experience these variations will depend on their size and mobility (Pincebourde & Woods, 2020).

Joint species distribution modelling allowed us to identify the specific effects of each environmental variable. From our hurdle approach, we could conclude that temperature and light affect communities in a two-step process. First, temperature acts as a filter determining which species from the local species pool are present within given conditions. Second, the species, once present, partition along a light intensity gradient. These results confirm that temperature can be an important factor limiting the activity of ectotherm pollinators (Clarke & Robert, 2018; Nielsen et al., 2017; Polatto et al., 2014), even for bumblebees that have developed thermoregulatory capabilities that enable them to be active in cold conditions (Lundberg, 1980). The lower relative importance of time of day might be due to the fact that bumblebees are able to forage across the entire day, from sunrise to sunset (Chapman et al., 2022; Hall et al., 2021). Our results emphasize the importance of understanding microhabitat niches for explaining species coexistence and for predicting their future responses to climate change, although the potential role of interspecific competition in shaping the observed communities still needs to be investigated directly (cf. Bowers, 1985). Our findings also highlight the important role that forested ecosystems could play as refugia for insect pollinators, something that is usually overlooked in pollination studies (Allen & Davies, 2023; Mola et al., 2021; Pincebourde & Woods, 2020; Ulyshen et al., 2023). In a climate warming context, tree canopy has the potential to buffer climate variation (Haesen et al., 2021), for example, by limiting the effects of climate warming on the thermophilization of plant communities (Zellweger et al., 2020) and acting as climate micro-refugia climate for ectotherms (Lenoir et al., 2017).

Our analyses showed that bumblebee species use microhabitats that differ in both light intensity and temperature. The responses of bumblebee species to these parameters were only slightly supported by the functional traits of body size and eye parameter, which might be due to the limited number (seven) of species/species groups observed in our study or that we only included data from queens. As such, we cannot draw strong conclusions about the specific relationships between traits and microhabitat characteristics. However, the three species with a high eye parameter (indicating an adaptation for foraging in dim light), *B. pratorum*, *B. pascuorum* and *B. hypnorum*, were associated with the low values of the light intensity gradient in the NMDS analysis (Figure S1). This is consistent with previous work in both diurnal bumblebees and tropical bees (Bartholomée et al., 2023; Streinzer et al., 2016; Tichit et al., in preparation). Further work on more diverse communities (*e.g.,* later in the season when more workers and more species are out foraging), with the inclusion of a higher number of functional traits involved in thermoregulation (*e.g.,* hairiness (Peters et al., 2016)) and visual traits (*e.g.,* diverse compound eye and ocelli traits) might help to identify specific relationships between functional traits and environmental variables.

## Conclusions

Our results show that bumblebee community composition depends on environmental conditions, with temperature determining what species are present and light intensity influencing the abundance of those species. The filtering role of temperature might indicate a stronger selection pressure when it comes to thermal niche when compared to light niche in diurnal pollinators, raising concerns for the consequences of climate warming on bumblebee and other insect pollinator communities. Our findings also pinpoint the central – but overlooked – role of microhabitat variations when studying the responses of ectotherms to climate. Our findings highlight that working towards a deeper and more exhaustive understanding of the relationships between pollinator and micro-climatic conditions is an urgent matter, particularly in forests, which may provide critical buffering zones that allow pollinators to mitigate the extreme effects of climate change.

## Acknowledgements

We thank Ola Langvall from SLU for helping with the determination of date of the bilberry blooming with his phenology model and Øystein Opedal for his support and guidance in the joint species distribution modelling. This work was part of the inter-disciplinary research environment project INVISMO funded by the Swedish Research Council (2018-06238). Additional funding was provided by a young researcher grant awarded to EB from the Swedish Research Council (2014-04762) and a young researcher grant from FORMAS awarded to OB (2023-01621). The analyses were performed with the support of a computer funded by Kungliga Fysiografiska Sällskapet i Lund, for Endowments for the Natural Sciences, Medicine and Technology – Biology (application 39016, year 2017). The research presented in this paper is a contribution to the Strategic Research Area “Biodiversity and Ecosystem Services in a Changing Climate”, BECC, funded by the Swedish government.

