## Supplementary material for "Warm or bright – temperature and light microhabitat use in insect pollinators": Table S1

**Table S1:** Traits value of the inter-tegular distance (ITD) and eye-parameter of all the observed bumblebee species and caste, as well as their source, adapted from Bartholomée et al. (2023).

| **Species*caste** | **Trait value (±SD) (*n =* sample size)** | **Comment** | **Source** |
| --- | --- | --- | --- |
| *B. hypnorum* queen | Eye parameter: 0. 726 (*n* = 1)*  ITD: 5.47 mm (±0.18) (*n* = 10) | The eye parameter is the value from the *B. hypnorum* worker. | Tichit *et al.* (Tichit et al., in preparation)  Romain Carrié, pers. com. |
| *B. lapidarius* queen | Eye parameter: 0.69 (*n* = 1)*  ITD: 5.58 mm (±0.16) (*n* = 10) | The eye parameter is the value from the *B. lapidarius* worker. | Tichit *et al.* (Tichit et al., in preparation)  Romain Carrié, pers. com. |
| *B. pascuorum* queen | Eye parameter: 0.734 (*n* = 1)*  ITD: 4.63 mm (±0.11) (*n* = 10) |  | Tichit *et al.* (Tichit et al., in preparation)  Romain Carrié, pers. com. |
| *B. pratorum* queen | Eye parameter: 0.828 (*n* = 1)*  ITD: 4.33 mm (±0.26) (*n* = 10) |  | Tichit *et al.* (Tichit et al., in preparation)  Romain Carrié, pers. com. |
| *B. soroeensis* queen | Eye parameter: 0.708 (*n* = 1)*  ITD: 4.71 mm (±0.18) (*n* = 10) | The eye parameter is the value from the *B. soroeensis* worker. | Tichit *et al.* (Tichit et al., in preparation)  Romain Carrié, pers. com. |
| *B. bohemicus* queen - cuckoo | Eye parameter: 0.597 (*n* = 1)*  ITD: 6.40 mm (*n* = 1)* | The eye parameter is the value of a queen of *B. bohemicus* standing for the cuckoo group. | Tichit *et al.* (Tichit et al., in preparation) |
| *B. terrestris* queen - BTC | Eye parameter: 0.688 (*n* = 1)*  ITD: 5.70 mm (±0.26) (*n* = 10) | The eye parameter is the value available of one queen *B. lucorum.* | Tichit *et al.* (Tichit et al., in preparation)  Romain Carrié, pers. com. |

*When only data for one individual was available, standard deviation is missing.

^†^It was not possible to compute the standard deviation, as only the mean and the sample size were available.

Tichit, P., Kendall, L., Olsson, P., Taylor, G., Bodey, A.J., Rau, C., Caplat, P., Smith, H.G. & Baird, E. (submitted) Country-wide shifts of visual communities along a habitat gradient. Do sensory traits drive community assembly in bumblebees?
