## Supplementary material for "Warm or bright – temperature and light microhabitat use in insect pollinators": Table S2

**Table S2:** Diagnostics of model convergence for the six models for both Beta and Gamma parameters: the mean effective sample size – with a good convergence indicated by a value close to the real number of samples (4000) and the mean and maximum potential scale reduction factors (PSRF), all close to one, indicating a good convergence.

| **Model** | **Parameter** | **Mean effective sample size** | **Mean PSRF** | **Max. PSRF** |
| --- | --- | --- | --- | --- |
| ***Occurrence*** | | | | |
| **M0** | **Beta** | **4061** | **1.0006** | **1.0023** |
|  | **Gamma** | **3811** | **1** | **1** |
| **M1** | **Beta** | **3899** | **1.0008** | **1.0029** |
|  | **Gamma** | **3957** | **1.0004** | **1.0012** |
| **M2** | **Beta** | **3568** | **1.0428** | **1.3518** |
|  | **Gamma** | **3934** | **1.0001** | **1.0012** |
| ***Conditional abundance*** | | | | |
| **M0** | **Beta** | **4081** | **1.0002** | **1.0005** |
|  | **Gamma** | **4000** | **1** | **1** |
| **M1** | **Beta** | **4114** | **1.0004** | **1.0024** |
|  | **Gamma** | **4087** | **1.0003** | **1.0007** |
| **M2** | **Beta** | **4040** | **1.0003** | **1.0014** |
|  | **Gamma** | **4057** | **1.0010** | **1.0038** |
