## Supplementary material for "Warm or bright – temperature and light microhabitat use in insect pollinators": Table S3

**Table S3:** Explanatory and predictive powers of the null models M0 for the occurrence and conditional abundance models – respectively measured by Tjurs’ R^2^ and R^2^. Partitioning of this explained variance in percentage by the different random effects (RE): plot identity (plot), sampling day (day) and time of day.

| **Occurrence model** | **Explanatory power**  **Tjurs’ R^2^** | **Predictive power**  **Tjurs’ R^2^** | **Proportion of the explained variance by the RE plot** | **Proportion of the explained variance by the RE day** | **Proportion of the explained variance by the RE time of day** |
| --- | --- | --- | --- | --- | --- |
| **Cuckoo** | 0.26 | 0.16 | 0.03 | 0.12 | 0.85 |
| ***B. hypnorum*** | 0.02 | -0.0004 | 0.36 | 0.28 | 0.36 |
| ***B. lapidarius*** | 0.18 | 0.12 | 0.06 | 0.74 | 0.20 |
| ***B. pascuorum*** | 0.07 | 0.01 | 0.80 | 0.04 | 0.16 |
| ***B. pratorum*** | 0.05 | 0.02 | 0.18 | 0.71 | 0.11 |
| ***B. soroeensis*** | 0.08 | 0.04 | 0.12 | 0.57 | 0.31 |
| **BTC** | 0.02 | -0.004 | 0.32 | 0.33 | 0.35 |
| **Conditional abundance model** | **Explanatory power**  **R^2^** | **Predictive power**  **R^2^** | **Proportion of the explained variance by the RE plot** | **Proportion of the explained variance by the RE day** | **Proportion of the explained variance by the RE time of day** |
| **Cuckoo** | 0.25 | 0.03 | 0.07 | 0.06 | 0.87 |
| ***B. hypnorum*** | 0.71 | 0.03 | 0.19 | 0.05 | 0.76 |
| ***B. lapidarius*** | 0.25 | 0.08 | 0.06 | 0.07 | 0.88 |
| ***B. pascuorum*** | 0.31 | 0.12 | 0.23 | 0.04 | 0.73 |
| ***B. pratorum*** | 0.17 | 0.004 | 0.58 | 0.11 | 0.30 |
| ***B. soroeensis*** | 0.37 | 0.21 | 0.05 | 0.08 | 0.87 |
| **BTC** | 0.57 | 0.41 | 0.13 | 0.28 | 0.59 |
