## Supplementary material for "Warm or bright – temperature and light microhabitat use in insect pollinators": Table S6

**Table S6.** Community response to the different environmental variables as supported by their functional traits (ITD and eye parameter) (Tjur’s R^2^ and R^2^ for occurrence and conditional abundance models, respectively) in M2.

| **M2** | **Occurrence model**  **(overall trait support: 0.15)** | **Conditional abundance model (overall trait support: 0.07)** |
| --- | --- | --- |
| **(Intercept)** | 0.44 | 0.29 |
| **Light intensity (linear term)** | 0.33 | 0.32 |
| **Light intensity (second order term)** | 0.29 | 0.33 |
| **Temperature (linear term)** | 0.59 | 0.27 |
| **Temperature (second order term)** | 0.42 | 0.20 |
| **Flower density** | 0.22 | 0.20 |
