## Supplementary material for "Warm or bright – temperature and light microhabitat use in insect pollinators": Table S7

**Table S7:** Explanatory and predictive powers of the models M2 with environmental variables and functional traits for the occurrence and conditional abundance models – respectively measured by Tjurs’ R^2^ and R^2^. Partitioning of this explained variance in percentage by the different explanatory variables: light intensity, temperature and flower density as well as random effects (RE): plot identity (plot), sampling day (day) and time of day. BTC stands for *Bombus terrestris* complex.

| **Occurrence model** | **Cuckoo** | ***B. hypnorum*** | ***B. lapidarius*** | ***B. pascuorum*** | ***B. pratorum*** | ***B. soroeensis*** | **BTC** |
| --- | --- | --- | --- | --- | --- | --- | --- |
| **Explanatory power Tjurs’ R^2^** | 0.29 | 0.07 | 0.23 | 0.16 | 0.10 | 0.15 | 0.10 |
| **Predictive power Tjurs’ R^2^** | 0.26 | 0.04 | 0.18 | 0.08 | 0.06 | 0.9 | -0.007 |
| **Light intensity** | 7.7 | 32.1 | 4.1 | 22.1 | 10.9 | 21.8 | 13.4 |
| **Temperature** | 78.9 | 22.0 | 32.9 | 16.8 | 9.7 | 23.7 | 29.0 |
| **Flower density** | 4.5 | 8.1 | 35.0 | 7.0 | 56.5 | 15.9 | 30.3 |
| **RE plot** | 1.6 | 8.2 | 3.1 | 36.1 | 7.4 | 3.7 | 10.1 |
| **RE day** | 2.5 | 4.8 | 20.9 | 13.1 | 8.6 | 28.0 | 3.8 |
| **RE time of day** | 4.8 | 24.8 | 4.0 | 4.8 | 7.0 | 6.9 | 13.4 |
| **Conditional abundance model** | **Cuckoo** | ***B. hypnorum*** | ***B. lapidarius*** | ***B. pascuorum*** | ***B. pratorum*** | ***B. soroeensis*** | **BTC** |
| **Explanatory power Tjurs’ R^2^** | 0.24 | 0.75 | 0.25 | 0.32 | 0.20 | 0.38 | 0.66 |
| **Predictive power Tjurs’ R^2^** | 0.12 | 0.52 | 0.17 | 0.12 | 0.01 | 0.11 | 0.56 |
| **Light intensity** | 38.4 | 42.6 | 57.0 | 46.1 | 14.5 | 41.8 | 9.6 |
| **Temperature** | 25.7 | 13.5 | 14.4 | 6.3 | 8.6 | 7.8 | 13.6 |
| **Flower density** | 9.7 | 8.4 | 7.3 | 7.5 | 8.7 | 7.9 | 18.6 |
| **RE plot** | 6.8 | 18.9 | 6.1 | 23.1 | 55.9 | 6.7 | 9.7 |
| **RE day** | 7.6 | 10.3 | 8.5 | 5.1 | 4.6 | 7.8 | 24.3 |
| **RE time of day** | 11.7 | 6.3 | 6.7 | 11.9 | 7.8 | 27.9 | 24.3 |
