## Supplementary material for "Warm or bright – temperature and light microhabitat use in insect pollinators": Table S8

**Table S8:** Species-level responses to environmental variable. Species occurrence in Model 2. Mean Beta parameters based on the posterior probability in the HMSC models 2, their credible intervals (CI) and their support. Values in grey correspond to a posterior support < 95%. BTC stands for *B. terrestris* complex.

| **Beta**  **[95% CI]**  **Support** | **(Intercept)** | **Light intensity (linear term)** | **Light intensity (second order term)** | **Temperature (linear term)** | **Temperature (second order term)** | **Flower density** |
| --- | --- | --- | --- | --- | --- | --- |
| **Cuckoo** | -2.294  [-10.094; -0.795]  0.99 | -0.029  [-1.118; 1.107]  0.52 | 0.011  [-0.061; 0.082]  0.62 | 0.499  [0.248; 0.766]  1.00 | -0.012  [-0.021; -0.004]  1.00 | 0.009  [-0.005; 0.024]  0.89 |
| ***B. hypnorum*** | 0.723  [-2.641; 4.293]  0.67 | -0.426  [-1.315; 0.408]  0.84 | 0.011  [-0.047; 0.072]  0.65 | 0.189  [-0.002; 0.383]  0.97 | -0.005  [-0.013; 0.001]  0.94 | -0.008  [-0.023; 0.007]  0.85 |
| ***B. lapidarius*** | -3.122  [-6.753; -0.043]  0.98 | 0.083  [-0.662; 0.947]  0.57 | -0.011  [-0.067; 0.039]  0.65 | 0.310  [0.147; 0.484]  1.00 | -0.009  [-0.015; -0.003]  1.00 | 0.029  [0.010; 0.047]  1.00 |
| ***B. pascuorum*** | 2.914  [-0.554; 6.393]  0.95 | -0.497  [-1.281; 0.319]  0.88 | 0.007  [-0.047; 0.052]  0.60 | 0.136  [-0.025; 0.298]  0.95 | -0.001  [-0.007, 0.005]  0.67 | -0.016  [-0.033; 0.000]  0.97 |
| ***B. pratorum*** | 2.282  [-1.478; 6.183]  0.88 | -0.371  [-1.337; 0.579]  0.77 | 0.020  [-0.042; 0.058]  0.73 | 0.040  [-0.138; 0.216]  0.67 | -0.001  [-0.008; 0.005]  0.66 | -0.023  [-0.036; -0.010]  1.00 |
| ***B. soroeensis*** | 0.834  [-2.533; 4.159]  0.69 | -0.510  [-1.354; 0.313]  0.89 | 0.017  [-0.037; 0.083]  0.73 | 0.251  [0.082; 0.431]  1.00 | -0.007  [-0.014; - 0.001]  0.99 | 0.016  [-0.001; 0.035]  0.96 |
| **BTC** | 10.890  [-7.641; 134.481]  0.60 | -4.763  [-55.577; 1.351]  0.64 | 0.373  [-0.098; 4.346]  0.62 | 0.184  [-3.170; 2.536]  0.84 | 0.014  [-0.032; 0.296]  0.28 | 0.467  [-0.014; 4.440]  0.90 |
