## Supplementary material for "Warm or bright – temperature and light microhabitat use in insect pollinators": Table S11

**Table S11:** Relationships between species functional traits and environmental variable. Species conditional abundance in M2. Mean Gamma parameters based on the posterior probability in the HMSC M2, their credible intervals (CI) and their support. Values in grey correspond to a posterior support < 95%.

| **Beta**  **[95% CI]**  **Support** | **(Intercept)** | **Light intensity (linear term)** | **Light intensity (second order term)** | **Temperature (linear term)** | **Temperature**  **(second order term)** | **Flower density** |
| --- | --- | --- | --- | --- | --- | --- |
| **(Intercept)** | 7.336  [-99.784; 112.251]  0.56 | -2.090  [-14.888; 11.383]  0.62 | -0.003  [-0.782; 0.831]  0.50 | 0.999  [-0.853; 4.448]  0.72 | -0.014  [-0.144; 0.117]  0.59 | 0.086  [-0.749; 0.886]  0.59 |
| **ITD** | -2.654  [-11.531; 6.339]  0.73 | 0.263  [-0.898; 1.412]  0.68 | -0.002  [-0.054; 0.076]  0.52 | 0.049  [-0.075; 0.344]  0.62 | -0.002  [-0.013; 0.009]  0.64 | 0.004  [-0.064; 0.077]  0.54 |
| **Eye parameter** | 9.012  [-82.074; 100.805]  0.58 | 0.631  [-11.089; 12.003]  0.55 | 0.027  [-0.585; 0.798]  0.553 | -1.453  [-0.736; 1.682]  0.82 | 0.028  [-0.087; 0.140]  0.69 | -0.147  [-0.849; 0.585]  0.67 |
