## Supplementary material for "Warm or bright – temperature and light microhabitat use in insect pollinators": Figure S1

**Complementary multivariate analyses**

**Drivers of bumblebee community composition**

1. **Methods**

As species richness was not significantly different between the observations taken at sunrise/sunset (for 1h) and during the rest of the day (15 min of observation; linear regression, *F*_1, 224_=1.05, *P* = 0.31), we converted community observations to the equivalent of 1 h observation time by multiplying the observations that were taken over 15 min (*i.e.,* those taken outside sunrise and sunset) by 4. To explore how environmental variables drive differences between communities, we ran non-metric multidimensional scaling (NMDS) analysis (e.g. as done in Dwyer et al., 2021) with the function ‘metaMDS’. We used the Bray-Curtis dissimilarity measure (designed for count data) (Borcard et al., 2011). NMDS is an unconstrained ordination based on species and not environmental data (as opposed to the RDA). For each of our four variables – light intensity, temperature, time, and flower density – we separately aligned the first dimension of the two-dimensional NMDS (‘MDSrotate’ function) with each environmental variable. The relationships were assessed using a Shepard diagram (function ‘stressplot’). For each environmental variable, we fitted a quadratic trend surface (‘ordisurf’ function) that we overlaid on the NMDS plot (note that the overlaid surfaces do not influence the ordination).

2. **Results**

The importance of light intensity, temperature, time and flower density on community composition was also supported by the NMDS analysis, which allowed us to explore the effect of each environmental variable separately. The stress of the two-dimension NMDS was 0.14, which shows a respect of dissimilarities (Clarke, 1993). The four environmental variables were significantly associated with bumblebee community composition with the trend surface of light intensity, temperature and time respectively explaining 20 %, 27 % and 22 % of the observed deviance (Figure S1A-C). In contrast, the trend surface of flower density explained 14 % of the observed variance (Figure S1D).

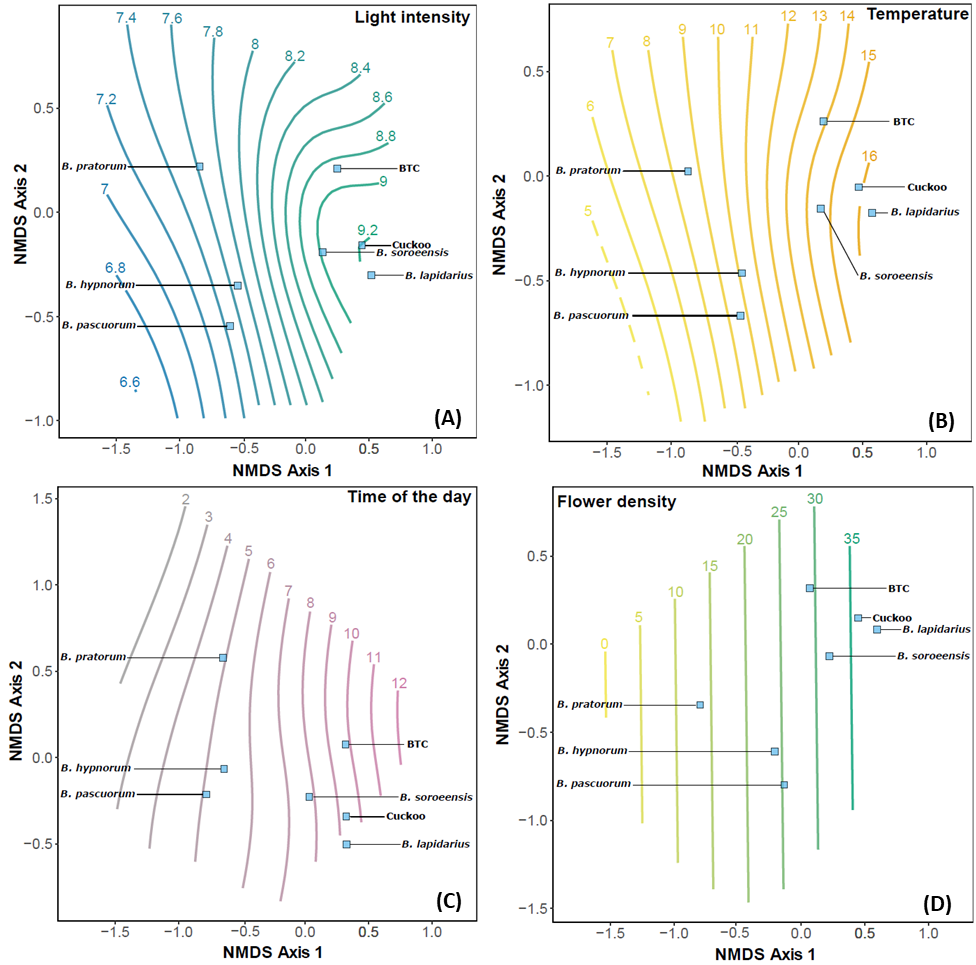

**FIGURE S1.** How bumblebee species segregate along environmental gradients, as analysed with non-metric dimensional scaling (NMDS). The trend surfaces of (A) light intensity (log-transformed lux, blue-to-green lines) (*F* = 13.36, e.d.f. = 3.26, *P* < 0.001, n = 226), (B) temperature (°C, yellow-to-orange lines) (*F* = 19.54, e.d.f. = 3.29, *P* < 0.001, n = 226), (C) time (hours from sunrise, grey-to-purple lines) (*F* = 15.15, e.d.f. = 3.05, *P* < 0.001, n = 226) and (D) flower density (number of flowers/0.25 m^2^, yellow-to-green lines) (*F* = 8.58, e.d.f. = 1.88, *P* < 0.001, n = 226) were overlaid on the species NMDS. Non-metric goodness-of-fit of the ordination: R^2^ = 0.98. Species are represented by blue squares. BTC stands for *B. terrestris* complex, cuckoo for the group of cuckoo species

Tichit, P., Kendall, L., Olsson, P., Taylor, G., Bodey, A.J., Rau, C., Caplat, P., Smith, H.G., & Baird, E. (in preparation) Country-wide shifts of visual communities along a habitat gradient. Do sensory traits drive community assembly in bumblebees? .
